# RDoC Mechanisms of Transdiagnostic Polygenic Risk for Trajectories of Depression: From Early Adolescence to Adulthood

**DOI:** 10.1101/2020.04.01.020495

**Authors:** James J. Li, Qi Zhang, Qiongshi Lu

**Affiliations:** Department of Psychology, University of Wisconsin-Madison; Waisman Center, University of Wisconsin-Madison; Department of Educational Psychology, University of Wisconsin-Madison; Department of Biostatistics and Medical Informatics, University of Wisconsin-Madison

**Keywords:** Depression, adolescence, RDoC, development, genetics

## Abstract

There is substantial heterogeneity in the development of depression across early adolescence into adulthood. Yet, little is known about the risk factors underlying individual differences in the development of depression. For instance, despite the discovery of genetic variants for depression, there is also significant genetic overlap between depression and other mental disorders. Thus, depression may have etiologically complex (i.e., transdiagnostic) origins when accounting for its heterogeneous developmental presentations. This study examined the association between a transdiagnostic polygenic score for psychopathology (*p*-factor PGS) and depressive trajectories, spanning early adolescence into adulthood, in the National Longitudinal Study of Adolescent to Adult Health. We also examined whether the Research Domains Criteria (RDoC) negative valence (i.e., negative emotionality), positive valence (i.e., novelty seeking), and cognitive systems (i.e., picture vocabulary) could explain how the *p*-factor PGS eventuates into the various pathways of depressive development. Four trajectories of depression were identified: *low depression* (78.9%)*, low increasing* (7.3%)*, high declining* (8.2%), and *early adult peaked* (5.7%). The *p*-factor PGS was only associated with the trajectory that showed increasing depression over time – *low increasing.* There was also a specific indirect effect by which the association of *p*-factor PGS on *early adult peaked* and *high declining* depression was partially mediated by negative emotionality, but not by picture vocabulary or novelty seeking. Our findings reinforce the crucial role of development in genetically-informed RDoC models of depression, as there appear to be distinct correlates and risk factors that underlie the various developmental pathways of depression. Clinical implications were also discussed.

**General Scientific Summary:** There are individual differences in how depression symptoms progress over time, but little is known about the risk factors that underlie these various patterns of development. This study suggests that there are distinct correlates and risk factors that underlie the various developmental pathways of depression. We found that transdiagnostic polygenic risks for psychopathology are directly associated with worsening patterns of adolescent to adult depression and indirectly associated with the less severe patterns of depression via negative emotionality.

---

Adolescence is a period of heightened vulnerability to depression (Hankin et al., 2015). As many as 50% of adolescents self-report clinically significant symptoms of depression (Kessler et al., 2001), and an estimated 8-20% of individuals will have a major depressive disorder before the age of 18 (Johnson et al., 2018). While depression generally declines from the middle adolescence into adulthood (Ge et al., 2001), there is substantial heterogeneity in how depression develops between individuals (Costello et al., 2008; Dekker et al., 2007; Olino et al., 2010). A better understanding of the processes that underlie this heterogeneity can reveal the subgroup of adolescents at greatest risk for negative outcomes and for whom prevention and/or interventions can be prioritized. However, relatively little is known about the risk factors that associate with individual differences in depressive development.

Prior prospective longitudinal studies of adolescent to adult trajectories of depression have identified distinct patterns during this epoch. Olino and colleagues (2010) identified 6 depression trajectories from the Oregon Adolescent Depression Project where only one trajectory had a high initial status and increasing depression over time; all other classes had consistently low initial rates of depression, where class differences were in their severity and directionality of slopes over time. Costello and colleagues (2008) examined depression symptoms in the National Longitudinal Study of Adolescent to Adult Health (Add Health) to identify 4 trajectories of adolescent (age 12) to adult (age 25) depression, where the classes reflected a *stable low depressed* group, and *early high declining depressed* group, and *a late escalating depressed* group. Notably, they also identified a single trajectory characterized by increasing levels of depression into adulthood (although they had a low initial status during adolescence). Overall, these studies shed important light on the various patterns of depressive development from adolescence to adulthood, including high risk individuals for whom depression worsens over time. However, identifying the etiological processes that underlie these developmental patterns remains a challenge.

Emerging discoveries in genomics seem poised to reveal the basic mechanisms underlying depression more broadly, especially considering the heritability of the disorder (37-48%; Corfield et al., 2017). The recent genome-wide association study (GWAS) on major depressive disorder (*N*=135,458 cases and 344,901 controls) revealed 44 genetic variants that were largely and unsurprisingly expressed in the brain (Wray et al., 2018). Still, these discoveries have yet to reveal any genetic or neurobiological mechanisms unique to depression. For instance, robust genetic correlations were observed between depression and every mental disorder assessed, including bipolar disorder (*r*=.32), schizophrenia (*r*=.34), and anxiety disorders (*r*=.80) (Wray et al., 2018). The depression-related *RBFOX1* gene is also enriched in the GWASs for autism and schizophrenia. Overall, the lack of etiologic clarity gained from GWAS thus far may reflect the high degree of heterogeneity (including developmental) across the mental disorders. It is possible that some patterns of development in depression may reflect more transdiagnostic and etiologically complex origins than others.

Additionally, these findings call into question the utility of identifying “genes for depression” when it appears that many of the genes identified across psychiatric GWASs overlap with one another. Recently developed methods in genetics can now characterize *transdiagnostic* genetic risks for psychopathology instead, reflecting the genetic covariation between multiple mental disorders that are correlated (Grotzinger et al., 2019; Selzam et al., 2018). This approach mirrors the clinically complexity of our patients, who frequently present with symptoms of multiple disorders. Thus, this study examines how transdiagnostic genetic risk factors eventuate into depression-specific pathways of development.

One framework that might elucidate *how* transdiagnostic genetic factors eventuate into the various depressive trajectories is the Research Domains Criteria (RDoC) project (Cuthbert, 2014). RDoC is a framework that aims to provide an integrative understanding of psychopathological mechanisms, from genes to neural circuits to behavior. Given the cardinal symptoms of depression, RDoC investigations of adolescent depression have naturally focused on measures within the negative valence (e.g., negative emotionality; Gore & Widiger, 2018; Woody & Gibb, 2015), positive valence (e.g., novelty seeking; Olino, 2016; Ortin et al., 2012), and cognitive systems (e.g., verbal reasoning and knowledge) (Goodall et al., 2018). However, developmentally-sensitive RDoC investigations are rare (Franklin et al., 2015) and few studies have examined the extent to which the RDoC subconstructs relate to trajectories of depression over time. Moreover, we are not aware of any studies have integrated a genetic methodology to interrogate these subconstructs within a prospective longitudinal framework.

This study integrates powerful methods in genetics with the RDoC framework to understand the mechanisms by which transdiagnostic genetic risks eventuate into specific developmental trajectories of depression. We examined longitudinal trajectories of depression across four Waves of Add Health, spanning ages 13 to 32. We then examined the association of the transdiagnostic genetic factor, measured as a *p*-factor polygenic score (PGS), along with RDoC measures within the negative valence, positive valence, and cognitive systems with the depression trajectories. This analysis was then followed by testing multiple mediation models that examined the direct and indirect effects by which the *p*-factor PGS was associated with individual differences in depression trajectories via the RDoC subconstructs.

## Method

### Participants

Add Health is a stratified sample of adolescents in grades 7-12 from high schools across the U.S. Data were collected from adolescents, parents, fellow students, school administrators, siblings, friends and romantic partners across four Waves: Wave I (1994-1995, grades 7-12, *N*=20,745), Wave II (1995-1996, grades 8-12, *N*=14,738), Wave III (2001-2002, ages 18-26, *N*=15,197), and Wave IV (2007-2008, ages 24-32, *N*=15,701). The current analyses were performed for the subset of Add Health individuals where both genotypic and phenotypic information were available (*N*=7,088). 46% of this sample was male, and the racial-ethnic composition was 63.6% Caucasian (including Hispanic), 20.7% African American, .2% Native American, 5.1% Asian, and 10.3% “Other.”

### Measures

#### Depression

Depression symptoms were measured using an abbreviated version of the Center for Epidemiologic Studies Depression Scale (CES-D) (Radloff, 1977) that were assessed across all four Waves of Add Health. Respondents provided responses, ranging from 0 (*never* or *barely*) to 3 (*most of the time* to *all of the time*), to items on several depression domains: somatic complaints (“bothered by things” or “can’t keep mind on tasks”), depressive affect (“felt depressed,” “unable to shake off the blues,” or “felt like life has been a failure”), positive affect (“felt as good as others” or “enjoyed life”), and interpersonal problems (“you feel disliked by people”). Total CES-D scores were computed. The internal consistencies of the depression items at Waves I-IV were above α=.80.

#### Cognitive System

The Add Health Picture Vocabulary Test (AHPVT) was administered to adolescents at Wave I to measure receptive vocabulary, verbal ability and scholastic aptitude. This measure is most aligned with the RDoC cognitive system subconstruct, *language*. For this test, the interviewer read a word aloud and the participant selected an illustration that best fit its meaning. Each word had four simple, black-and-white illustrations arranged in a multiple-choice format. The standardized AHPVT score was used.

#### Negative Valence System

While negative emotionality is notably absent from RDoC, it is highly correlated with enumerated subconstructs within the negative valence system (Gore & Widiger, 2018). We used 6 factor-analytically derived items from Wave I to assess negative emotionality as described in Young and Beaujean (2011). Respondents were asked to rate on how much they agreed or disagreed, from 1 (*strongly agree*) to 5 (*strongly disagree*), with statements that were aligned with neuroticism on the NEO Personality Inventory (Costa & McCrae, 1992) (e.g., NEO: “have a low opinion of myself,” Add Health: “you have a lot of good qualities”). The measure had good internal consistency, α=.86.

#### Positive Valence System

We examined the novelty seeking subscale for the measure of self-control as described in Beaver et al. (2009). Novelty seeking is most aligned with the positive valence system subconstruct, *reward responsivity*. The novelty seeking subscale was comprised of 7 items assessed during Wave III (items related to novelty seeking were not assessed at earlier Waves). Respondents were asked to rate on how true statements were to them, ranging from 1 (*not true*) to 5 (*very true*). Example items include “I often try new things just for fun and thrills, even if most people think they are a waste of time” and “when nothing new is happening, I usually start looking for something exciting”. The internal consistency of this scale was strong, α=.84.

### Genotyping and Polygenic Score Computation

Saliva were obtained from participants at Wave IV. Genotyping was done on the Omni1-Quad BeadChip and the Omni2.5-Quad BeadChip. Add Health European genetic samples were imputed on Release 1 of the Human Reference Consortium (HRS r1.1). Non-European samples were imputed using the 1000 Genomes Phase 3 reference panel. To derive a *p*-factor PGS, we modeled the multivariate genetic architecture of multiple traits from their GWAS summary statistics using Genomic Structural Equation Modeling (*GenomicSEM*) in *R* (Grotzinger et al., 2019). Briefly, the *p*-factor PGS was derived from the joint analysis of five genetically correlated psychiatric case-control GWAS – major depressive disorder, schizophrenia, bipolar disorder, post-traumatic stress disorder, and anxiety disorders (information on each GWAS is available through the Psychiatric Genomics Consortium). We used a GWAS *p*-value threshold of 1.0 to incorporate all available SNP information as well as to minimize bias due to overfitting.

### Statistical Analyses

#### Step 1

Latent class growth (LCG) models of depression were fit in Mplus 7.4 (Muthén & Muthén, 2015). Data were restructured such that time was represented by age. We used the full Add Health sample for this analysis. The form factor of the LGC model was identified by comparing class fit statistics across four different models with the following parameters: 1) intercept only, 2) intercept and slope, 3) intercept, slope, and quadratic, and 4) intercept, slope, quadratic, and cubic. We report the Akaike Information Criterion (AIC) and Bayesian Information Criterion (BIC) values (lower values indicate better fit to the data), entropy (values approaching 1.0 indicate clear delineation of classes), and the Lo-Mendell-Rubin likelihood ratio *p*-value test (low *p*-value rejects the k-1 class model in favor of the k class model). Optimal model identification was based on the evaluation of these fit statistics as well as their interpretability from a substantive and theoretical perspective.

#### Step 2

A multinomial logistic regression was also performed in Mplus where depression classes (identified in Step 1) were regressed on the RDoC variables (i.e., negative emotionality, picture vocabulary, novelty seeking) and *p*-factor PGS while also controlling for sex, age, parental education, and genetic ancestry based on the first 10 principal components (PCs) of the genetic data. We controlled for sex because of several lines of evidence that indicated that the trajectories of depression from early adolescence to young adulthood were invariant by sex (Costello et al., 2008; McLaughlin & King, 2015). Relative risk ratios (RRR) were estimated, reflecting the association between each variable and their contributions to the relative risk of membership into a depression trajectory relative to a reference class. For the current analysis, the class representing the greatest number of individuals (usually a normal or low depression class) will serve as the reference group. Add Health survey weights for incorporated into all models.

#### Step 3

Multiple mediation models were performed in *R version 3.6.1*. In multiple mediation, the total effect, *c*, reflects the pathway of *p*-factor PGS to the risk of being in a depression class (relative to the reference class) independent of the effects of the RDoC mediators – negative emotionality, picture vocabulary, and novelty seeking, *a*_*i*_ reflects the effect of *p*-factor PGS on each mediator, and *b*_*i*_ reflects the effect of each mediator on the risk of being in the depression class relative to the reference class. The direct effect, *c’*, reflects the association of *p*-factor PGS on the risk of being in a depression class after accounting for the mediators. Specific indirect effects of the *p*-factor PGS predicting depression class membership via RDoC measures were also estimated. Models controlled for sex, age, parent education, and 10 genetic PCs.

## Results

### Depression Symptom Trajectories

Supplemental Table 1 provides fit statistics for each LGC model. Based on model fit indices and considerations of the model’s substantive interpretability (i.e., alignment with prior findings and developmental theories of depression), the 4-class LGC model of depression was optimal (see Figure 1). The *low depression* class (78.9% of individuals) exhibited consistently low levels of depression symptoms over time. The *low increasing depression* class (7.3%) had a similarly low levels of depression at baseline (age 13), but exhibited a steady increase in depression beginning in the early 20’s. The *high declining depression* class (8.2%) had a high initial status of depression in early adolescence but had a steep decline in depression during the teenage years, culminating in virtually indistinguishable levels of depression relative to the *low depression* class by the mid-20’s. Finally, the *early adult peak depression* class (5.7%) was characterized by low levels of depression at baseline that steadily increased up until ~age 23, followed by a steady decline in depression *to low depression* levels after the age 23 inflection point.

**Figure 1.**
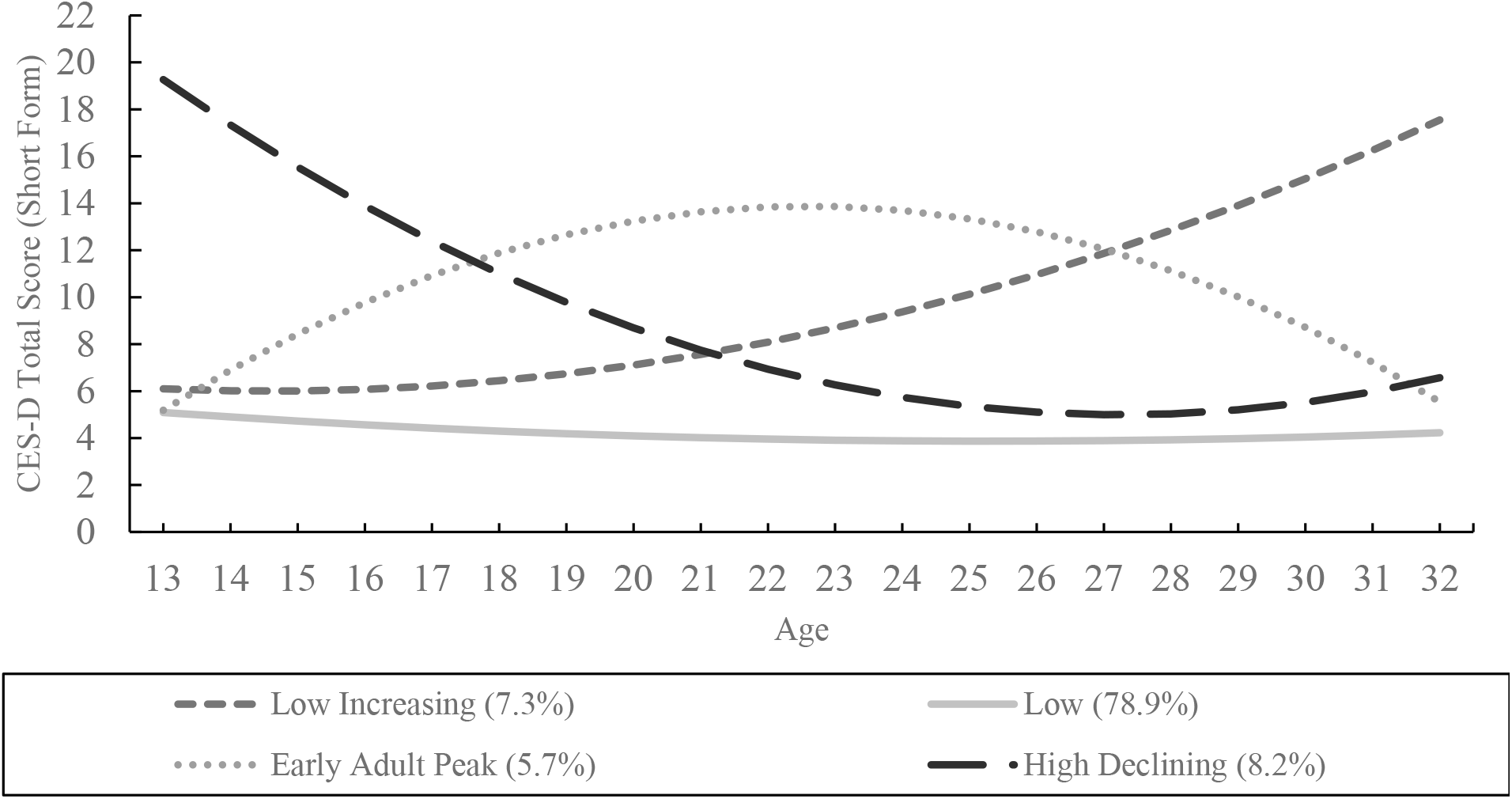
Latent class growth trajectories depression

### Multinomial Logistic Regression of Depression Symptom Trajectories

Supplemental Table 2 provides the results from a multinomial logistic regression that examined the association of each RDoC measure and the *p*-factor PGS on the relative risk of belonging to each depression class relative to the reference class (i.e., the *low depression* class).

For *the low increasing depression* class, higher negative emotionality (RRR=1.121, *p*<.01, 95% CI=1.064, 1.180), novelty seeking (RRR=1.058, *p*<.01, 95% CI=1.032, 1.085), and the *p*-factor PGS (RRR=1.371, *p*<.01, 95% CI=1.072, 1.753), but not picture vocabulary, increased one’s relative risk of belonging into this class relative to the *low depression* class.

For the *early adult peak depression* class, higher negative emotionality (RRR=1.203, *p*<.01, 95% CI=1.123, 1.288), lower picture vocabulary scores (RRR=.984, *p*<.01, 95% CI=.974, .995), and higher novelty seeking (RRR=1.092, *p*<.01, 95% CI=1.039, 1.126), but not *p*-factor PGS were associated with belonging to this class relative to the *low depression* class.

Finally, for the *high declining depression* class, only higher negative emotionality (RRR=1.362, *p*<.01, 95% CI=1.306, 1.420) and lower picture vocabulary (RRR=.986, *p*<.01, 95% CI=.979, .994) were associated with belonging to this class relative to the *low depression* class; neither novelty seeking nor *p*-factor PGS were associated with *high declining depression* class membership.

### Multiple Mediation Analysis

Figure 2a-c shows the indirect effects of the *p*-factor PGS on each depression class through the simultaneous effects of the RDoC mediators relative to the *low depression* class. Supplemental Table 3 provides the total and indirect specific effects.

**Figure 2.**
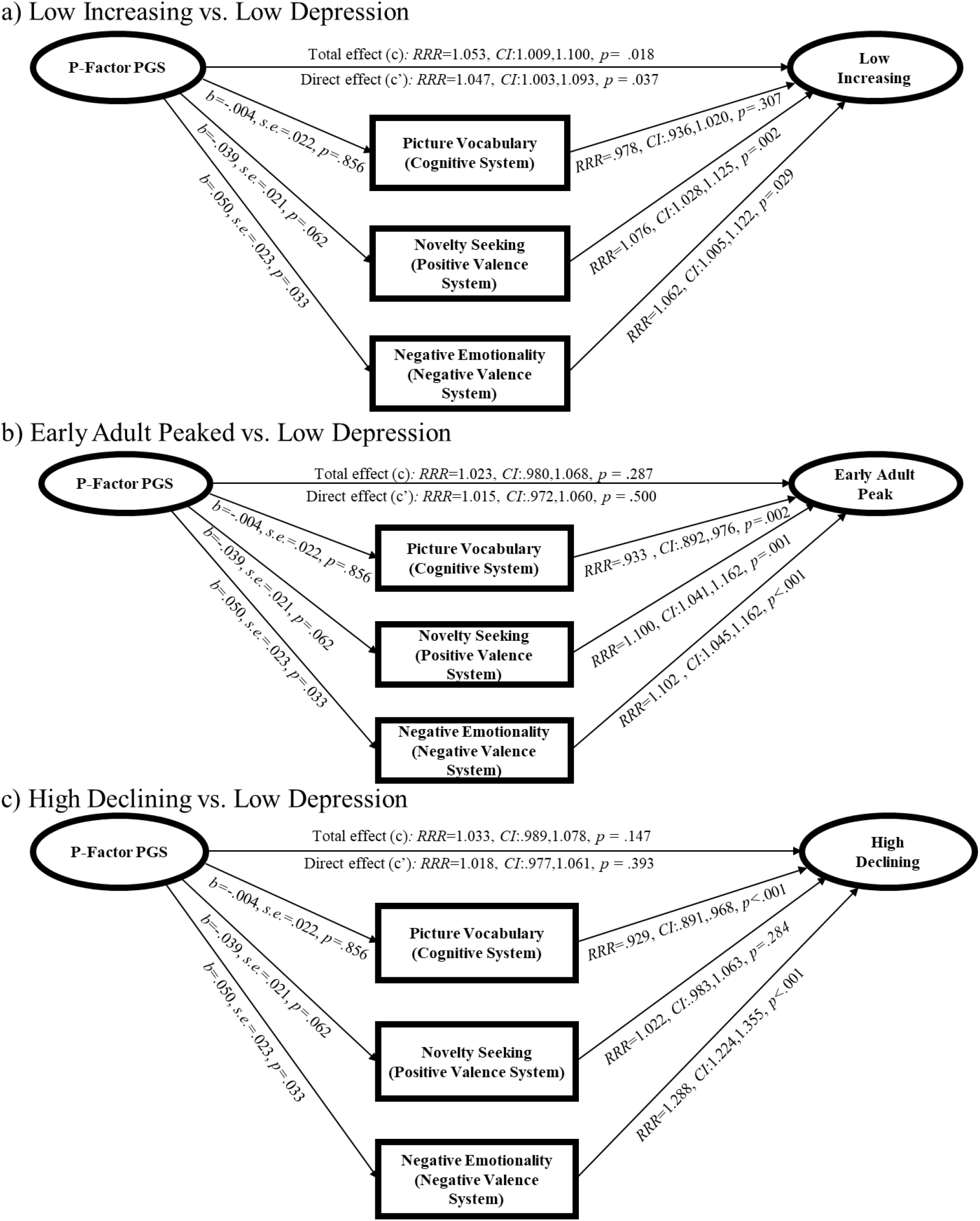
Direct and indirect effects of the *p*-factor PGS on developmental trajectories of depression via RDoC measures

For the *low increasing depression* model (Figure 2a), higher *p*-factor PGS increased one’s risk of *low increasing depression* class membership by 5.3% (i.e., the total effect, without accounting for the mediators) relative to *low depression class* membership (95% CI=1.009, 1.100). After accounting for the RDoC mediators (i.e., the direct effect), higher *p*-factor PGS increased one’s risk of *low increasing depression* class membership by 4.7% relative to *low depression* class membership (95% CI=1.003, 1.096). However, the total indirect effect of *p*-factor PGS on this trajectory was not significant (95% CI=0, .001) and no specific indirect effect via RDoC mediators emerged.

For the *early adult peak depression* model (Figure 2b), neither the total nor the direct effect of *p*-factor PGS on class membership was significant. However, there was a specific indirect effect of *p*-factor PGS on early *adult peak depression* class membership via negative emotionality (95% CI=.001, .010).

Similarly, for the *high declining depression* model (Figure 2c), the total and the direct effect of p-factor PGS on class membership were not significant. However, there was a specific indirect effect of *p*-factor PGS via negative emotionality (95% CI=.001, .024).

## Discussion

We identified 4 distinct, early adolescent (age 13) to adult (age 32) trajectories of depression (i.e., *low depression, low increasing, high declining*, and *early adult peak*) that were generally consistent with findings from previous prospective longitudinal studies (Costello et al., 2008; Dekker et al., 2007; Olino et al., 2010). Notably, we found that the *p*-factor PGS was only associated with the trajectory that did not exhibit begin to exhibit depression until adulthood – the *low increasing* depression trajectory. This finding seems contrary to two recent genetically-informed studies of depression that found greater depression-specific polygenic liability underlying earlier onset depression relative to later onset depression (Power et al., 2017; Rice et al., 2019), which also fits with findings that individuals with early-onset depression are typically at higher risk for chronic depression and maladjustment in later life (Hammen et al., 2008). However, we did not identify a developmentally persistent subtype of depression, as the *low increasing* trajectory was the only one characterized by increasing depression over time. We surmise that this group’s worsening depression during adulthood may be reflective of their greater overall genetic liability. That is, greater genetic liability to psychopathology may be associated with other psychiatric comorbidities that are more prominent during adulthood (e.g., substance use), whereas depression-specific polygenic liability may be more associated with the earlier-onset and possibly more “pure” forms of depression (*a la* Power et al., 2017; Rice et al., 2019). These hypotheses should be tested in future prospective longitudinal studies.

Next, we found that RDoC measures were differentially associated with the depression trajectories, suggesting unique cognitive, behavioral, and emotional processes underlying the different patterns of depressive development. While it was not surprising that negative emotionality (reflecting RDoC negative valence system) was broadly associated with the depression trajectories, we observed a specific indirect effect by which the association of *p*-factor PGS on *early adult peak* and *high declining* depression was partially mediated by negative emotionality, but not by picture vocabulary (cognitive system) or novelty seeking (positive valence system). This finding was supported by prior evidence that negative emotionality may be a mechanistic pathway through which early risk factors can lead to the development of depression (Barrocas & Hankin, 2011). Our results suggest that this mechanism may be specific to the patterns of depression that attenuate by or during the adult years (age ~30). Furthermore, while picture vocabulary was negatively associated with the *early adult peaked* and *high declining* depression trajectories as expected, we did not expect to find a positive association between novelty seeking with *low increasing* and *early adult peak* depression. High novelty seeking is a known risk factor for suicide and substance use among depressed adolescents (Ortin et al., 2012), but it was unclear why novelty seeking was more associated with the adult trajectories of depression specifically. Future developmental studies of depression should account for co-occurring substance use and suicide behavior to better understand the association of positive valence constructs and depression.

A few study limitations should be noted. First, depression and two of the RDoC measures (i.e., negative emotionality and novelty seeking) were assessed via self-report. Shared method variance could have inflated the associations between the RDoC measures (mediators) and depression (outcome). Additionally, while we focused on RDoC measures that were measured at the earliest timepoint (thus coinciding with the earliest timepoint of our depression trajectories), novelty seeking was only assessed at Wave III. This precluded our ability to fully temporally separate the mediators from the outcomes. Furthermore, while we accounted for population stratification by controlling for the first 10 genetic PCs, we are aware that these controls do not adequately account for the Eurocentric-bias in discovery GWASs (including for depression), which attenuates the polygenic prediction signal for non-Eurocentric target populations such as Add Health (Martin et al., 2019). Future investigations should anticipate the greater genetic prediction error that comes with using a European-based GWAS in admixed samples. Finally, and as mentioned throughout this discussion, we did not assess other behaviors that are known to covary with depression, such as anxiety, suicide, and substance use behaviors. Future studies should identify the risk factors for more complex developmental models of depression that also account of co-occurring conditions.

The RDoC framework promises to elucidate the behavioral and neurobiological mechanisms by which our genes eventuate into normal and abnormal behaviors, including psychopathology (Cuthbert, 2014). Our findings reinforce the importance of also accounting for development in RDoC models (Franklin et al., 2015), as there appear to be distinct correlates and risk factors that underlie the various developmental pathways of depression (Costello et al., 2008; Dekker et al., 2007; Olino et al., 2010). Adopting a developmental lens in our studies may also have important clinical implications. Characterizing developmental patterns of depression may help to identify the subgroup of adolescents who are at highest risk for negative outcomes in later life, and for whom RDoC-targeted interventions may be most effective. Individuals at higher risk may even benefit more from interventions relative to those at moderate and even low risk (Bakermans-Kranenburg & IJzendoorn, 2015). Considering their treatment implications, these hypotheses are worth exploring further through a developmental lens.

## Acknowledgements

This study was supported in part by a core grant to the Waisman Center from the *Eunice Kennedy Shriver* National Institute of Child Health and Human Development (U54 HD090256).

