## Supplemental Tables for "RDoC Mechanisms of Transdiagnostic Polygenic Risk for Trajectories of Depression: From Early Adolescence to Adulthood"

*Supplemental Table 1*. Fit statistics for the latent class growth analysis of depression

| | AIC | BIC | Entropy | LRT *p*-value |
|---|---|---|---|---|
| Intercept only models | | | | |
| 2 class | 342751.33 | 342939.67 | .78 | <.01 |
| 3 class | 342049.90 | 342253.94 | .74 | <.01 |
| 4 class | 342053.90 | 342273.63 | .76 | .50 |
| 5 class | 342057.90 | 342293.33 | .55 | .50 |
| 6 class | 342061.90 | 342313.02 | .71 | .50 |
| Intercept + Slope models | | | | |
| 2 class | 341147.15 | 341366.88 | .81 | <.01 |
| 3 class | 339263.58 | 339506.86 | .79 | <.01 |
| 4 class | 338649.85 | 338916.67 | .77 | .013 |
| 5 class | 338097.64 | 338388.00 | .77 | <.01 |
| 6 class | 337714.37 | 338028.28 | .73 | .19 |
| Intercept + Slope + Quadratic models | | | | |
| 2 class | 340936.53 | 341195.50 | .81 | <.01 |
| 3 class | 338915.78 | 339206.14 | .79 | <.01 |
| 4 class | 337998.48 | 338320.24 | .80 | .24 |
| 5 class | 337811.16 | 338164.31 | .82 | .69 |
| 6 class | 337242.52 | 337627.05 | .79 | .44 |
| Intercept + Slope + Quadratic + Cubic models | | | | |
| No convergence | | | | |

*Notes*. AIC= Akaike information criterion; BIC= Bayesian information criterion; LRT=Lo-Mendell-Rubin likelihood ratio test

*Supplemental Table 2*. Multinomial logistic regression predicting depression growth trajectories

| Variables | RRR | Std. Err. | *p*-value | 95% Confidence Interval | |
|---|---|---|---|---|---|
| | | | | Lower | Upper |
| LOW INCREASING DEPRESSION CLASS (7.3%) | | | | | |
| Sex | 1.825 | .300 | .000 | 1.317 | 2.529 |
| Age | .947 | .048 | .283 | .857 | 1.046 |
| Parental Education | .921 | .044 | .089 | .837 | 1.013 |
| Negative Emotionality | 1.121 | .029 | .000 | 1.064 | 1.180 |
| Picture Vocabulary | .995 | .003 | .083 | .989 | 1.001 |
| Novelty Seeking | 1.058 | .014 | .000 | 1.032 | 1.085 |
| *p*-factor PGS | 1.371 | .170 | .012 | 1.072 | 1.753 |
| EARLY ADULT PEAK DEPRESSION CLASS (5.7%) | | | | | |
| Sex | 2.640 | .676 | .000 | 1.590 | 4.383 |
| Age | 1.011 | .069 | .867 | .884 | 1.157 |
| Parental Education | .923 | .045 | .105 | .838 | 1.017 |
| Negative Emotionality | 1.203 | .042 | .000 | 1.123 | 1.288 |
| Picture Vocabulary | .984 | .005 | .004 | .974 | .995 |
| Novelty Seeking | 1.082 | .022 | .000 | 1.039 | 1.126 |
| *p*-factor PGS | 1.166 | .174 | .305 | .868 | 1.566 |
| HIGH DECLINING DEPRESSION CLASS (8.2%) | | | | | |
| Sex | 1.672 | .337 | .012 | 1.121 | 2.493 |
| Age | 1.323 | .062 | .000 | 1.205 | 1.453 |
| Parental Education | .999 | .045 | .982 | .913 | 1.093 |

| | | | | | |
|---|---|---|---|---|---|
| Negative Emotionality | 1.362 | .029 | .000 | 1.306 | 1.420 |
| Picture Vocabulary | .986 | .004 | .001 | .979 | .994 |
| Novelty Seeking | 1.021 | .014 | .141 | .993 | 1.050 |
| *p*-factor PGS | 1.143 | .102 | .137 | .958 | 1.365 |

*Note*. Genetic PCs 1-10 were included in the model but are not listed in the table above. These results are available upon request.

*Supplemental Table 3*. Total and specific indirect effects of the *p*-factor PGS on depression class membership via RDoC mediators

| | | | | 95% Confidence Interval | |
|---|---|---|---|---|---|
| Class contrasts | Indirect Effect | Effect | Std. Err. | Lower | Upper |
| Increasing Depression vs. Low Depression | Total | .006 | .003 | .000 | .011 |
| | Picture Vocabulary | .000 | .001 | -.001 | .001 |
| | Novelty Seeking | .003 | .002 | -.001 | .006 |
| | Negative Emotionality | .003 | .002 | -.001 | .007 |
| Early Adult Peak Depression vs. Low Depression | Total | .009 | .004 | .000 | .016 |
| | Picture Vocabulary | .000 | .002 | -.003 | .003 |
| | Novelty Seeking | .004 | .002 | -.001 | .008 |
| | Negative Emotionality | .005 | .002 | .001 | .010 |
| High Declining Depression vs. Low Depression | Total | .014 | .006 | .001 | .026 |
| | Picture Vocabulary | .000 | .002 | -.003 | .003 |
| | Novelty Seeking | .001 | .001 | -.001 | .003 |
| | Negative Emotionality | .013 | .006 | .001 | .024 |
